# Intrinsic Hippocampal-Caudate Interaction Correlates with Human Navigation

**DOI:** 10.1101/116129

**Authors:** Xiang-Zhen Kong, Yi Pu, Xu Wang, Shan Xu, Xin Hao, Zonglei Zhen, Jia Liu

## Abstract

It has been indicated that both egocentric and allocentric representation systems exist in parallel, and combine to support spatial navigation according to the task. Identifying the neuronal mechanisms and functional roles of the interactions between the two systems promises to provide new insights into the organization of human navigation network. Here we combined resting-state fMRI and behavioral tasks to investigate how the core structures of these systems (i.e., hippocampus and caudate) functionally interact, and further examined their behavioral relevance in navigation in healthy young adults (N = 190). We found a slightly positive connectivity between the hippocampus and caudate (especially in good navigators), suggesting an effective hippocampal-caudate cross talk, which may facilitate the functional integration of the two systems. Interestingly, the hippocampal-caudate interaction correlated with better self-reported navigation ability. Moreover, using an extra 3D pointing task in virtual reality, we found that individual’s behavioral performance could be largely predicted by the hippocampal-caudate interactions. Overall, our study demonstrated the intrinsic interaction between two representation systems of the navigation network and its functional roles in behaviors, and further study on dynamic interaction would help us understand better their role in normal aging and psychiatric disorders.

## Introduction

To navigate the world is crucial for human in everyday life. It has been indicated that in our brain both egocentric and allocentric representation systems exist in parallel, and combine to support spatial memory according to the task (Burgess N 2006). The allocentric representations allow navigation from new starting locations (Hartley T et al. 2003) based on the large-scale environmental configuration (Iaria G et al. 2003; Doeller CF et al. 2008) or recognition of places from a new viewpoint (King JA et al. 2002; Lambrey S et al. 2008), while the egocentric representations allow navigation via a fixed route (Hartley T *et al.* 2003; Iaria G *et al.* 2003) or relative to a single landmark (Doeller CF *et al.* 2008). Although there are considerable individual preferences in use of one or the other of these representations, it is expected that, during real-world navigation, switching between them adaptively would facilitate accurate navigation (Andersen RA 1997; Bremmer F et al. 2001). Actually, emerging behavioral and neuroimaging evidence has demonstrated the cooperative nature of interactions between two systems of spatial representation underlying navigation (Voermans NC et al. 2004; Brown TI et al. 2012; Rice JP et al. 2015). For example, in patients with Huntington’s disease, the hippocampus compensates for gradual caudate nucleus dysfunction with a gradual activity increase during route recognition to maintaining normal behavior (Voermans NC *et al.* 2004), while the hippocampus (O'Keefe J and L Nadel 1978; Morris RG et al. 1982) and the caudate nucleus (Cook D and RP Kesner 1988; Brasted PJ et al. 1997) have been indicated to be core structure in the two systems, respectively. Although a cooperative integration between two systems has been suggested, it is still unclear what the neural basis of this interaction is, and how the functional interaction supports human navigation behaviors.

Resting-state functional magnetic resonance imaging (fMRI) combined with functional connectivity analysis provides a unique method for examining the functional interactions among brain areas and their roles in behavioral variability. It is suggested that positive functional connectivity between two brain areas may represent an efficient inter-regional cross talk, which may facilitate certain functional processes. For instance, our recent study has shown that there is positive functional connectivity between two core brain areas in the face network (i.e., the occipital face area and fusiform face area), the strength of which predicts individual variability in face recognition ability (Zhu Q et al. 2011). Within the navigation network, for another instance, functional connectivity between the parahippocampal gyrus and hippocampus has been shown to be positively related to participants’ self-reported navigation ability (Wegman J and G Janzen 2011). In addition, resting state fMRI has been used for charting the navigation network in the human brain, which provides new insights into understanding the underlying organization of the functional network and the individual differences (Kong XZ, X Wang, et al. 2017). These suggest that, by examining the functional connectivity, we would be able to reveal the nature of functional interactions between regions and their functional roles in certain behaviors.

Here we combined resting-state fMRI and multiple behavioral tasks, including both selfreport and behavioral tasks, to investigate how core structures of the egocentric and allocentric representation systems (i.e., hippocampus and caudate) functionally interact, and further examined their behavioral relevance in navigation in healthy young adults (N = 190). We hypothesized that there would be a positive functional connectivity between the hippocampus and caudate, especially in good navigators. Moreover, we hypothesized that the hippocampal-caudate interactions could predict individual differences in navigation behaviors.

## Materials and Methods

### Participants

One hundred and ninety college students (117 females; mean age = 20.3 years, standard deviation (SD) = 0.91 years) from Beijing Normal University (BNU), Beijing, China, participated in the study. The dataset is part of the Brain Activity Atlas Project (BAA, http://www.brainactivityatlas.org/) (Kong XZ et al. 2014; Zhen Z et al. 2015; Kong XZ, Y Song, et al. 2017; Zhen Z et al. 2017). All participants had normal or corrected-to-normal vision. The study was approved by the Institutional Review Board of BNU.

Written informed consent was obtained from all participants before they took part in the experiment.

All participants (N = 190) underwent the resting-state fMRI scanning and no participant was excluded due to excessive head motion (2 mm in translation or 2 degree in rotation from the first volume in any axis) or visually detected registration errors (Zhen Z *et al.* 2015). In addition, most of these participants (N = 167; 104 females; mean age = 20.2 years, SD = 0.90 years) participated four behavioral assessments, including a standard questionnaire on general navigation ability in daily life, a computer test on small-scale spatial ability, a Raven task for general ability, and a 3D pointing task in a virtual environment task on a specific navigation performance. Seven participants (2 females) failed to complete the 3D pointing task and were therefore excluded from the corresponding analyses.

### MRI scanning

Scanning was conducted at BNU Imaging Center for Brain Research, Beijing, China, on a Siemens 3T scanner (MAGENTOM Trio, a Tim system) with a 12-channel phased-array head coil. Participants were instructed to relax without engaging in any specific task and to remain still with their eyes closed during the scan. The resting-state scan lasted 8 min and consisted of 240 contiguous echo-planar-imaging (EPI) volumes (TR=2000 ms; TE=30 ms; flip angle=90°; number of slices=33; matrix=64 × 64; FOV=200 × 200 mm^2^; acquisition voxel size=3.125 × 3.125 × 3.6 mm^3^).

High-resolution T1-weighted images were acquired with magnetization prepared gradient echo sequence (MPRAGE: TR/TE/TI=2530/3.39/1100 ms; flip angle=7^o^; matrix= 256 × 256) for spatial registration. One hundred and twenty-eight contiguous sagittal slices were obtained with 1 × 1 mm^2^ in-plane resolution and 1.33-mm slice thickness.

### Imaging data analysis: data preprocessing

For each participant, image preprocessing was performed with FMRIB Software Library (FSL, http://www.fmrib.ox.ac.uk/fsl/). Preprocessing included head motion correction (by aligning each volume to the middle volume of the image with MCFLIRT), spatial smoothing (with a Gaussian kernel of 6-mm full-width half-maximum), intensity normalization, and removal of linear trend. Next, a temporal band-pass filter (0.01–0.1 Hz) was applied with *fslmaths* to reduce low frequency drifts and high-frequency noise.

Registration of each participant’s high-resolution anatomical image to a common stereotaxic space (the Montreal Neurological Institute (MNI) 152-brain template with a resolution of 2 × 2 × 2 mm^3^, MNI152) was accomplished using a two-step process. Firstly, a 12-degrees-of-freedom linear affine was carried out with FLIRT (Jenkinson M and S Smith 2001; Jenkinson M et al. 2002). Second, the registration was further refined with FNIRT nonlinear registration (Andersson JLR et al. 2007). Registration of each participant’s functional images to the high-resolution anatomical images was carried out with FLIRT to produce a 6-degrees-of-freedom affine transformation matrix.

To eliminate physiological noise, such as fluctuations caused by motion or cardiac and respiratory cycles, nuisance signals were regressed out using the methods described in previous studies (Fox MD et al. 2005; Biswal BB et al. 2010). Nuisance regressors included averaged cerebrospinal fluid signal, averaged white matter signal, global signal averaged across the whole brain, six head realignment parameters obtained by rigid-body head motion correction, and the derivatives of each of these signals. The 4-D residual time series obtained after removing the nuisance covariates were registered to MNI152 standard space by applying the previously calculated transformation matrix.

### Imaging data analysis: quality control

Two predefined criteria were used to access the qualities of functional MR images. First, to minimize the artifacts due to motion during scanning, the participants with excessive head motion (greater than 2.0 ^o^ or 2.0 mm throughout the rs-fMRI scan) would be excluded from the further analysis. Second, to minimize the error caused by misalignment of functional and anatomical volumes, the participants with large error of registration would be excluded. Specifically, the registration quality was checked visually by overlaying the normalized volume on the MNI152 template. No participant showed excessive head motion or visually detectable registration errors; that is, no participant was excluded and the data from all participants were included in the following analyses.

### Imaging data analysis: ROI selection

Anatomical ROIs of the hippocampus and caudate (in each hemisphere) were derived using the Harvard-Oxford probabilistic atlas that was included with FSL. We defined our ROIs to only include voxels that had 50% or higher probability of being as corresponding anatomical label (left hippocampus: 4016 mm^3^; right hippocampus: 4248 mm^3^; left caudate: 3328 mm^3^; right caudate: 3672 mm^3^; Fig. 1). All subsequent analyses were performed separately for these ROIs.

**Fig. 1.**
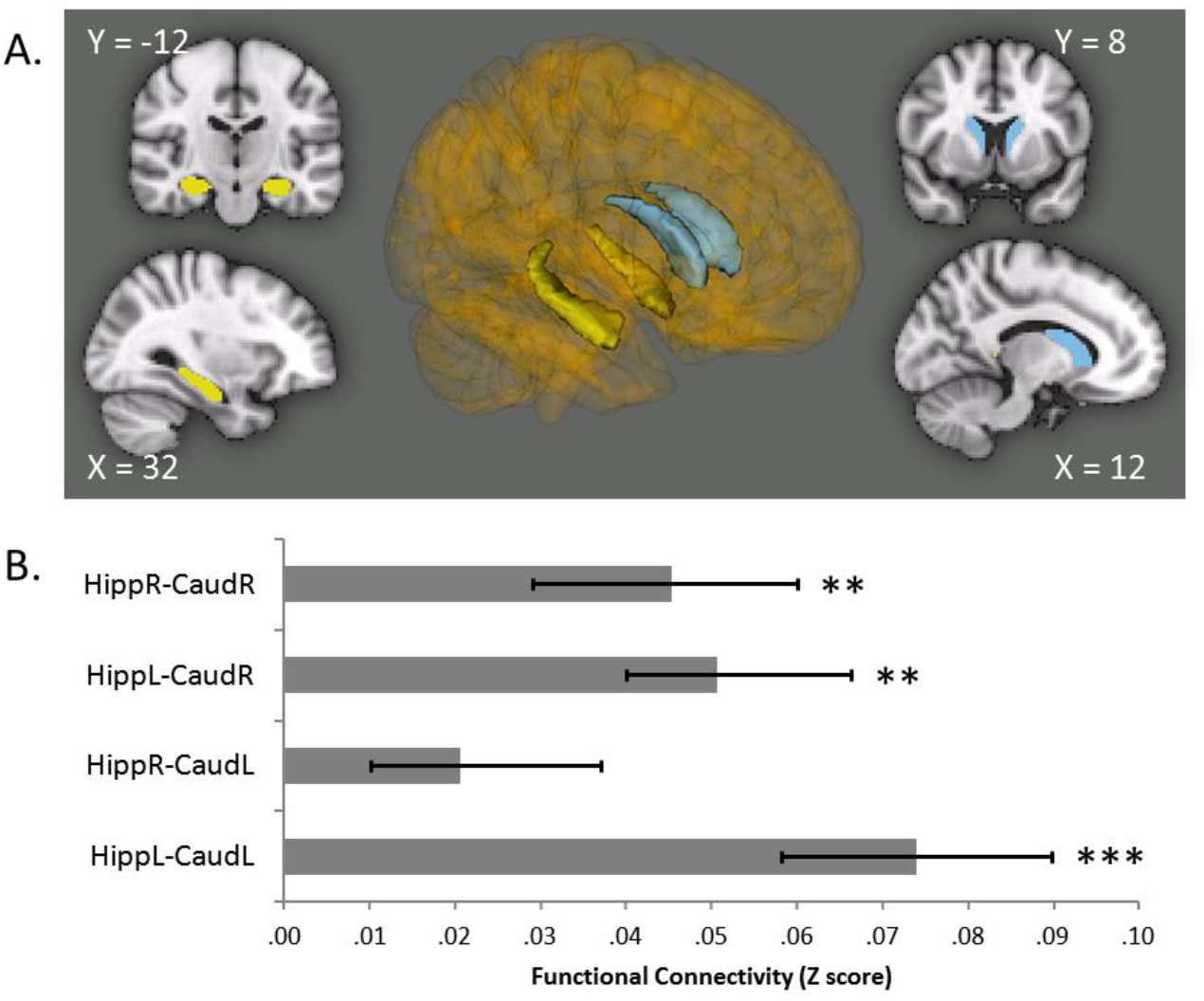
Functional connectivity between the hippocampus and caudate. (A) Anatomical ROIs of the hippocampus and caudate (in each hemisphere). Yellow: Hippocampus; Blue: Caudate. (B) Functional connectivity between the hippocampus and caudate in both ipsilateral and contralateral hemispheres revealed with resting-state fMRI. Error bars represent ± standard error of the mean (SEM). HippR-CaudR / HippL-CaudL, functional connectivity between the hippocampus and caudate in the same hemispheres; HippR-CaudL/ HippL_CaudR, functional connectivity between the hippocampus and caudate in the different hemispheres; R, right; L, left. ** p < 0.01, *** p < 0.001.

### Imaging data analysis: resting-state functional connectivity

After the preprocessing, a continuous time course for each ROI was extracted by averaging the time courses (from resting-state fMRI; 240 TRs; TR = 2000 ms) of all voxels in each of that ROI. Thus, we obtained a time course consisting of 236 data points (removed the first 4 data points) for each ROI and for each participant. Temporal correlation coefficients between the extracted time course from a given ROI and those from other ROIs were calculated to determine which regions were functionally correlated at rest. Correlation coefficients (r) were transformed to Gaussian-distributed z scores via Fisher’s transformation to improve normality, and these z scores were then used for further analyses (Fox MD et al. 2006).

### Behavioral tests: sense of direction scale

Navigational ability was operationalized as scores on the Santa Barbara Sense of Direction scale (SBSOD) (Hegarty M et al. 2002), which is a standard test on sense of direction in a large-scale environment, increasingly used as a reliable proxy for actual navigation ability (Janzen G et al. 2008; Wegman J and G Janzen 2011). SBSOD consists of 15 items. Example items are “I very easily get lost in a new city” and “I can usually remember a new route after I have traveled it only once”.

Participants were instructed to indicate the extent to which they agreed or disagreed with each statement in a 5-point Likert-type scale. The total score was used to index one’s navigation ability, with higher scores indicating better performance in daily navigation.

Note that previous studies have shown that people have explicit and accurate knowledge on their own navigation ability (Kozlowski LT and KJ Bryant 1977; Sholl MJ 1988; Wolbers T and M Hegarty 2010), and therefore it is not surprising that the scale, which is based on navigation experiences in daily life, has been found highly reliable (test-retest reliability: 0.91). Another reason of choosing the SBSOD is that it can be easily administrated and thus has been widely used as a reliable proxy for real-world navigation performance in a variety of neuroimaging studies (Epstein RA et al. 2005; Janzen G *et al.* 2008; Wegman J and G Janzen 2011; Auger SD et al. 2012; Wegman J et al. 2014). For instance, with structural MRI and DTI data, Wegman et al. (2014) have demonstrated that the gray and white matter of the caudate nucleus and medial temporal regions correlates with navigation ability measured by the SBSOD. With task fMRI, the strength of fMRI adaptation effect in the PPA correlates with SBSOD score (Epstein RA *et al.* 2005), and the effect of memory consolidation of landmarks in the hippocampus is observed only in good navigators who are screened by the SBSOD (Janzen G *et al.* 2008), whereas poor navigators who are also screened by the SBSOD are less reliable at identifying landmarks with reduced activation in the RSC and anterodorsal thalamus (Auger SD *et al.* 2012). With resting-state fMRI, Wegman and Janzen (2011) have demonstrated that the functional connectivity at rest between the PHG and the hippocampus/caudate is related to participants’ navigational ability measured by the SBSOD. Taken together, the SBSOD is valid to be used as a proxy of real-world navigation performance in neuroimaging studies.

### Behavioral tests: Raven’s advanced progressive matrices (RAPM)

To eliminate the possible influence of the general ability on the relationship between navigation ability and the intrinsic interaction connectivity of interest, individual’s general intelligence was measured using the standard RAPM (Raven J 1995). The number of correct responses to the test items of RAPM was used to index intelligence for this study.

### Behavioral tests: mental rotation task (MRT)

Mental rotation is a small-scale spatial ability, and considered relying on cognitive mechanisms different from spatial navigation. To investigate the navigation-specific nature of the observed brain-behavior association, we also measured individual’s small-scale spatial ability. Participants were administered the MRT (Shepard RN and J Metzler 1971), consisting of 40 trials. Each trial started with a blank screen for 0.5 s, followed by the first cube stimulus presented at the center of the screen. The three-dimensional asymmetrical assemblages of cube image were presented for 0.7s. After an ISI of 0.5 s, the second stimulus appeared for the same duration as the first one, with the viewpoint being changed. Subjects were instructed to indicate whether the second stimulus was the first one rotated or another stimulus as quickly as possible. Participants were given 3 min to finish all 40 trials, including 20 trials for ‘rotated’ condition and 20 for ‘another’ condition. The accuracy was used to index individual’s mental rotation ability.

### Behavioral tests: 3D pointing task

An objective measure of navigation ability was measured with a 3D pointing task in virtual mazes (Lawton CA and KA Morrin 1999). Mazes were created using Unity3D, which was used to present and control the experiment as well. Each maze consisted of a series of hallways of equal length and uniform texture and color (Fig. 4A, 4B). There was only one path that could be taken through each maze. No landmarks were present in the hallways, except for two words ‘Start’ and ‘Finish’ on the start and end of the pathway respectively. The point of view while moving through the mazes was approximately eye-level.

Mazes consisted of either one turn, two turns, four turns, or six turns. All turns were right angles (90^o^). There were two variations of the one-turn maze (either a right-hand turn or left-hand turn), which were used as practice mazes, and four variations of each of the two-, four-, and six-turn mazes used in experimental trials. These mazes were configured such that there were no more than two consecutive turns in the same direction (i.e., no loop).

There were a total of 24 mazes, consisting of a set of 12 mazes repeated twice but in a different order. Each set of 12 mazes contained four blocks of mazes, each block consisting of a different two-turn, four-turn, and six-turn maze, randomly ordered.

Participants were first given practice on the computer pointing task using the two one-turn mazes. Movement through the mazes was controlled by the computer with same speed (5 s per maze hallway). Participants controlled view direction at the end of the mazes using the computer mouse to point the start point. In practice trials, feedback would be given about the accuracy of their responses. Then, participants continued with the 24 experimental mazes and were given no feedback on these mazes. Participants were asked not to draw out a map of the mazes. Pointing error scores were computed as the absolute difference between the participant’s answer and the correct answer, with error scores up to 180^o^ possible in either direction.

To rule out the possible influence of video game playing, participants were also asked to rate the frequency with which they have played the video games requiring navigation through an environment on a 5-point scale, ranging from *Not at All Frequently* to *Very Frequently.*

### Statistical analyses: Hippocampal-caudate functional connectivity correlates with spatial navigation ability

To investigate the association between the intrinsic hippocampal-caudate interaction and navigation ability, we performed partial correlation analyses between navigation ability measured by SBSOD and the functional connectivity between the hippocampus and caudate, with age and sex controlled. Spearman correlation analysis was also performed to confirm the correlations observed. In this study, we investigated both the ipsilateral and contralateral connectivity between the hippocampus and caudate, including the connectivity between left hippocampus and left caudate (HippL-CaudL), connectivity between right hippocampus and right caudate (HippR-CaudR), connectivity between left hippocampus and right caudate (HippL-CauR), and connectivity between right hippocampus and left caudate (HippR-CaudL). False Discovery Rate (FDR) correction for multiple comparisons (as described in (Huang Y et al. 2013), see also (Hastie T et al. 2009)) was used to control the false positive rate. Associations with a corrected *p* value of < 0.05 were considered statistically significant.

Further, we investigated the directionality (i.e., positive or negative nature) of the resting-state functional connectivity between hippocampal and caudate in good and poor navigators. We first divided the participants into good (N = 81) and poor (N = 86) navigator groups using a mediation split (median SBSOD = 47), and then conducted one-sample t-test analysis on the hippocampal-caudate functional connectivity separately for each group.

### Statistical analyses: seed-based functional connectivity between hippocampal subfields and caudate correlates with navigation ability

Given both the hippocampus and caudate are anatomically and functionally heterogeneous (Gilbert PE et al. 2001; Yassa MA and CE Stark 2011; Robinson JL et al. 2012), besides the hippocampal-caudate functional connectivity analyses above, we further conducted seed-based voxel-wise functional connectivity analyses to investigate the observed brain-behavior association. Specifically, the hippocampus subfields, including CA1, CA2/3, CA4/DG, presubiculum, subiculum, fimbria and the hippocampal fissure, were defined using a new automated procedure (Van Leemput K et al. 2009) based on individual’s T1-weighted MRI data. Segmentation results were visually inspected for errors in all datasets, and no manual edit was done. Since the procedure was based on a probabilistic model, a 50% threshold was used to define the hippocampal subfield ROIs to ensure that there was no overlap between the ROIs. Then, a continuous time course for each ROI was extracted by averaging the time courses of all voxels in each of that ROI. For each voxel within the caudate, the functional connectivity with each hippocampal subfield ROI was calculated with a Pearson correlation analysis followed by a Fisher’s transformation. We conducted voxel-based multiple regression with SBSOD score as the independent variable. The confounding variables (i.e., age and sex) were controlled as mentioned above. We used a stringent threshold of *p* < 0.05 family-wise error (FEW) corrected for multiple comparisons.

### Statistical analyses: Hippocampal-caudate functional connectivity predicts individual’s performance in the 3D pointing task

To investigate whether the hippocampal-caudate functional connectivity at rest could predict individual’s performance in specific navigationally-relevant tasks, we conducted a 3D pointing task, in which participants were asked to point to the starting point of the route just taken at the end of a maze (see *Behavioral tests).* Initially, to replicate the brain-behavior association found with SBSOD, partial correlation analyses were performed between pointing error and the resting-state functional connectivity between the hippocampus and caudate, with age, sex and game playing frequency. The game playing frequency was controlled to rule out the possible influence of video game playing on the 3D task performance. Given that we has a priori hypotheses about the direction of the correlation (as shown below), we used one-tail significance threshold *p* < 0.05. Moreover, to estimate to what extent these hippocampal-caudate interactions together could predict individual’s performance in the pointing task, we used support vector regression (SVR) and leave-one-out cross-validation (LOOCV) procedure. The Pearson correlation coefficient between observed performance and the predicted score was calculated to provide measure of prediction accuracy.

## Results

### Resting-state interaction between the hippocampus and the caudate

We found that the hippocampus showed slightly positive functional connectivity with the caudate in both hemispheres (HippL-CaudL: 0.08 ± 0.017, t(189) = 4.66, *p* < 0.001; HippR-CaudR: 0.05 ± 0.015, t(189) = 3.01, *p* = 0.003) (Fig. 1B). The positive functional connectivity suggested cooperative functional interactions between the two structures (though the strength was relatively weak; see below for more discussion). Although the present study mainly focused on the hippocampal-caudate interactions in ipsilateral hemisphere, we included the hippocampal-caudate interactions between contralateral regions in this study. We found the left hippocampus showed positive functional connectivity with the right caudate (HippL-CauR: 0.05 ± 0.016, t(189) = 3.20, *p* = 0.002), while the functional connectivity between the right hippocampus and left caudate only showed a weak positive trend but no significant difference from zeros (HippR-CaudL: 0.02 ± 0.017, t(189) = 1.30, *p* = 0.20) (Fig. 1B). No sex difference was found *(ps* > 0.50).

### Hippocampal-caudate interaction correlates with navigation ability

Next, we investigated whether individuals with higher hippocampal-caudate connectivity at rest reflected individual variability in navigation ability.

To link the intrinsic hippocampal-caudate interaction to human navigation, we measured individual’s general navigation ability with the SBSOD. As expected, navigation score showed wide variability (from 27 to 74; 48.72 ± 9.55) across participants (Kong XZ, X Wang, *et al.* 2017). Consistent with previous studies (Astur RS et al. 1998; Moffata SD et al. 1998; Kong XZ, Y Huang, et al. 2017), males showed significantly higher scores than females (t(165) = 2.69; *p* = 0.008), suggesting better navigation ability in males on average. Further, we conducted correlation analyses between navigation ability and the functional connectivity between hippocampus and caudate, controlling for sex and age. Results showed that both ipsilateral and contralateral hippocampal-caudate interaction showed significantly positive correlation with individual’s navigation ability (Fig. 2; HippL-CaudL: *r* = 0.25, *p* = 0.001; HippR-CaudR: *r* = 0.19, *p* = 0.014; HippL-CaudR: *r* = 0.22, *p* = 0.004; HippR-CaudL: *r* = 0.24, *p* = 0.002). These results were confirmed with Spearman correlation analysis (HippL-CaudL: *rho* = 0.26, *p* = 0.001; HippR-CaudR: *rho* = 0.17, *p* = 0.027; HippL-CaudR: *rho* = 0.23, *p* = 0.003; HippR-CaudL: *rho* = 0.26, *p* = 0.001). That is, individuals with higher hippocampal-caudate interaction tend to possess better navigation ability, suggesting that the hippocampal-caudate interaction at rest would benefit individual’s daily navigation.

**Fig. 2.**
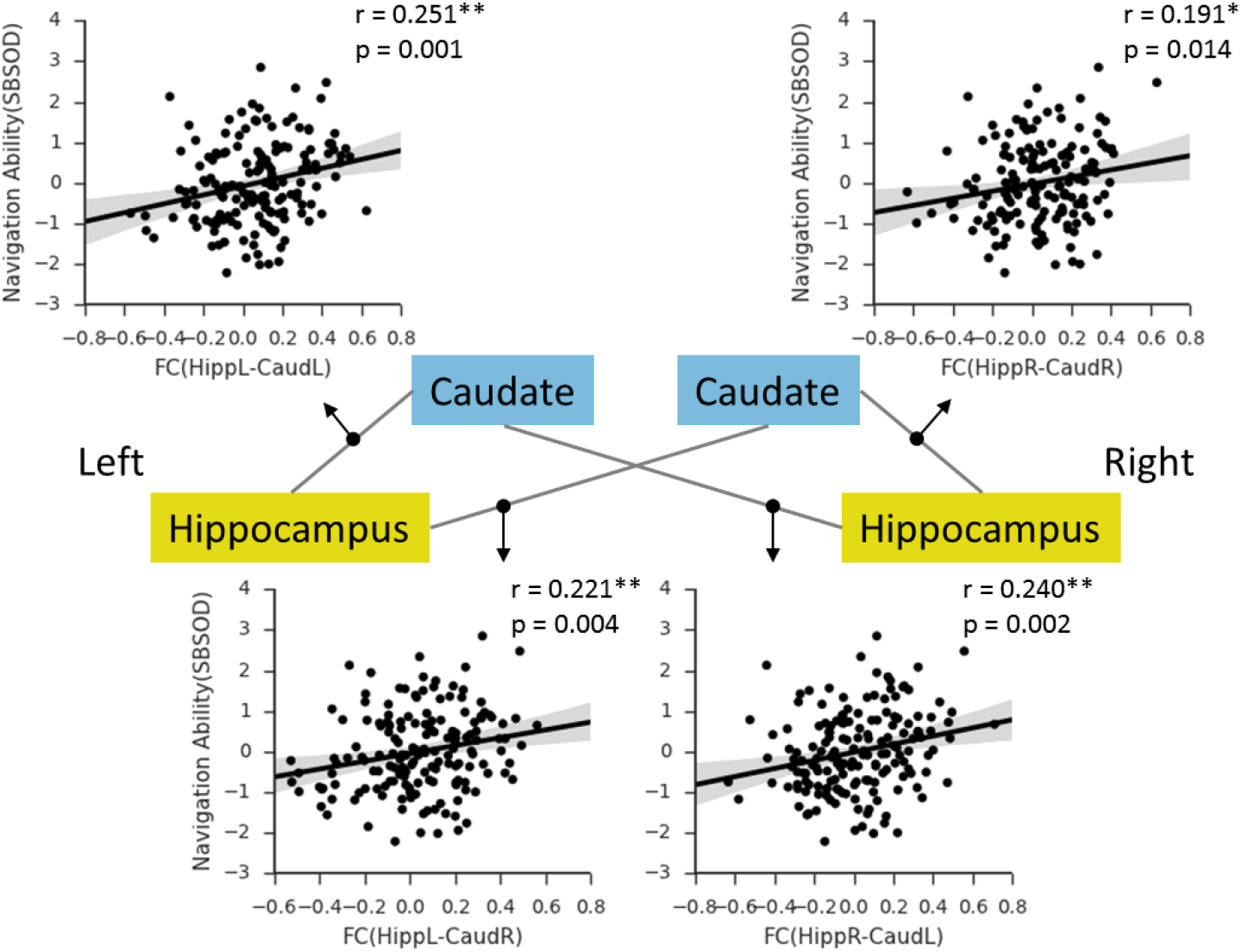
Correlations between the hippocampal-caudate functional connectivity and navigation ability measured with SBSOD. Shaded regions depict 95% confidence intervals. HippR-CaudR/ HippL-CaudL, functional connectivity between the hippocampus and caudate in the same hemispheres; HippR-CaudL/ HippL_CaudR, functional connectivity between the hippocampus and caudate in the different hemispheres; R, right; L, left. ** p < 0.001; *** p < 0.001.

Based on these findings, we further evaluated the nature of directionality of the functional connectivity by dividing the subjects into good (N = 81) and poor (N = 86) navigator groups using a mediation split (median SBSOD = 47). Results showed that the good navigator group, as expected, had positive functional connectivity between the hippocampus and the ipsilateral caudate (Hipp_L-Caud_L: t(80) = 5.56, *p* < 0.001; Hipp_R-Caud_R: t(80) = 3.58, *p* = 0.001), whereas the poor navigator group did not show the relationship (Hipp_L-Caud_L: t(85) = 0.46, *p* = 0.65; Hipp_R-Caud_R: t(85) = −0.35, *p* = 0.731) (Fig. 3). Similarly, the good navigator group showed positive functional connectivity between the hippocampus and the contralateral caudate (Hipp_L-Caud_R: t(80) = 5.11, *p* < 0.001; Hipp_R-Caud_L: t(80) = 2.75, *p* = 0.007), whereas the poor navigator group did not show the relationship (Hipp_L-Caud_R: t(85) = −0.64, *p* = 0.525; Hipp_R-Caud_L: t(85) = −0.17, *p* = 0.098) (Fig. 3). Hence, the positive interaction between the hippocampus and caudate was more prominent in good navigators, which suggests a positive link between the hippocampal-caudate interaction and navigation ability.

**Fig. 3.**
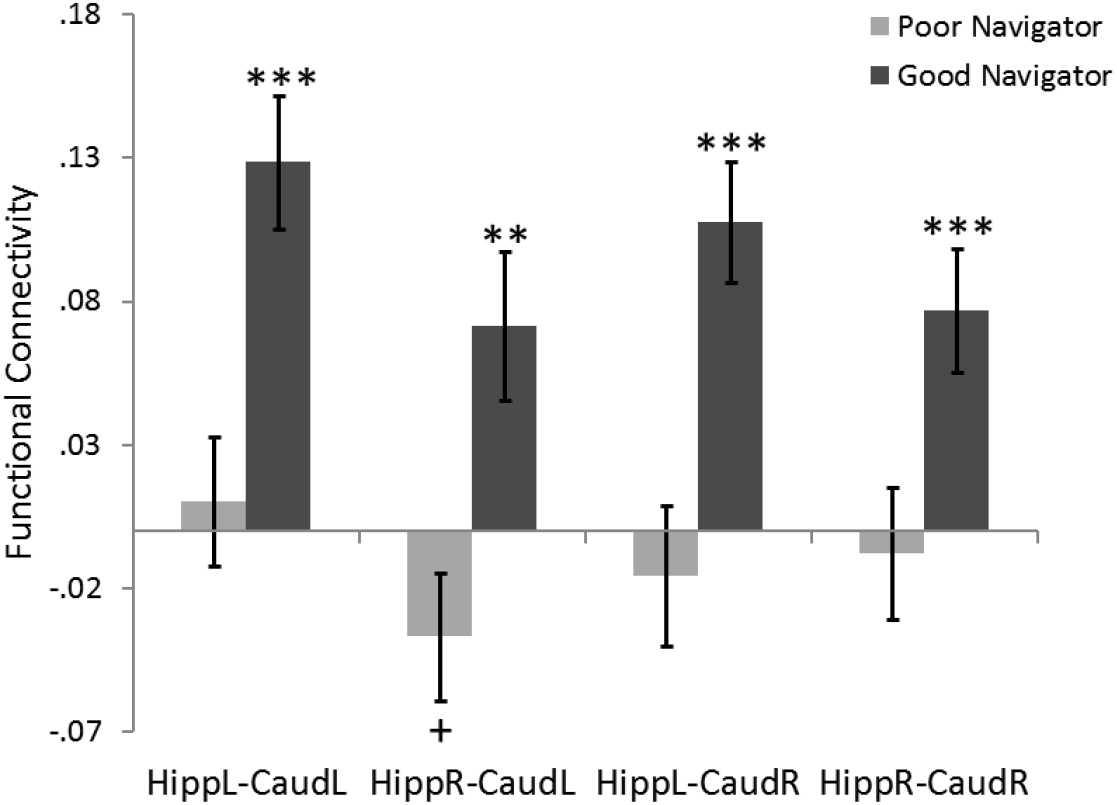
Functional connectivity between the hippocampus and caudate in good navigators and poor navigators. HippR-CaudR/ HippL-CaudL, functional connectivity between the hippocampus and caudate in the same hemispheres; HippR-CaudL/ HippL_CaudR, functional connectivity between the hippocampus and caudate in the different hemispheres; R, right; L, left. + p < 0.10; ** p < 0.01; *** p < 0.001.

### Navigation-specific nature of the findings

The hippocampus and caudate have been implicated as the cores of two systems of spatial representation underlying navigation. Next, we examined whether the behavioral correlates of the hippocampal-caudate interactions are specific to navigation. First, we asked whether the hippocampal-caudate interactions were related to non-navigation performance, including general ability and small-scale spatial ability. To rule out the possibility that general ability contributed to the observed brain-behavior associations, we measured individual’s general intelligence using the standard RAPM (Raven J 1995). We found that when the general intelligence was controlled, the observed associations remained significant (HippL-CaudL: *r* = 0.25, *p* = 0.002; HippR-CaudR: *r* = 0.19, *p* = 0.015; HippL-CaudR: *r* = 0.22, *p* = 0.004; HippR-CaudL: *r* = 0.23, *p* = 0.003), suggesting the observed associations were independent to general ability. Moreover, to further investigate the specificity of the observed associations, we measured individual’s small-scale spatial ability with MRT and found no significant correlations between small-scale spatial ability and the hippocampal-caudate interactions in the HippR-CaudR (r = 0.09, *p* = 0.230), HippL-CaudL (r = 0.13, *p* = 0.088) and the HippR-CaudL (r =0.06, *p* = 0.412). Although interaction in the HippL-CaudR showed a significant correlation with small-scale spatial ability (r = 0.17, *p* = 0.023), when the small-scale ability was controlled, the observed associations between navigation ability and hippocampal-caudate interactions remained significant (r = 0.21, *p* = 0.006). In sum, these findings suggest that the associations are specific to large-scale spatial ability (i.e, navigation ability).

Furthermore, as a control experiment for neural specificity, we conducted similar correlation analysis with functional connectivity between contralateral regions for the hippocampus and the caudate separately. We found that functional connectivity between contralateral analogs for either the hippocampus (r = 0.02, *p* = 0.757) or caudate (r = −0.09, *p* = 0.246) showed no correlation with navigation ability, suggesting the neural specificity of the observed associations between navigation and the hippocampal-caudate interaction.

Finally, given both the hippocampus and caudate are anatomically and functionally heterogeneous (Gilbert PE *et al.* 2001; Yassa MA and CE Stark 2011; Robinson JL *et al.* 2012), we further conducted seed-based functional connectivity analyses with hippocampal subfields as seeds. These seeds, including CA1, CA2/3, CA4/DG, presubiculum, subiculum, fimbria and the hippocampal fissure (Fig. 4A), were defined using a new automated procedure (Van Leemput K *et al.* 2009) based on individual’s T1-weighted MRI data. The results showed that resting-state functional connectivity of all hippocampal subfields with the caudate showed similar correlation with navigation ability, and in caudate it was mainly the dorsal medial caudate contributing to this association (Fig. 5).

**Fig. 4.**
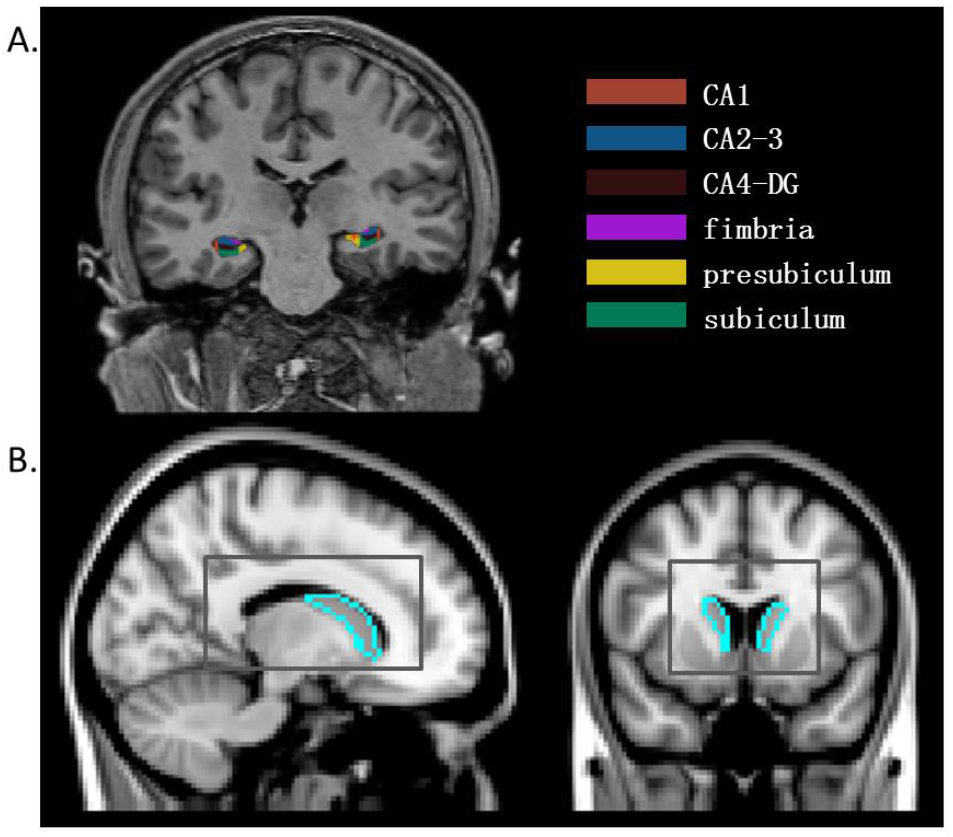
Anatomical ROIs of hippocampal subfields and caudate for seed-based analysis. (A) The hippocampus segmentation for one subject superimposed on the subject's T1-weighted scan in coronal view. (B). Region boundary of the ROI of caudate (blue) in sagittal and axial views, respectively and the region reference for Fig. 5 (gray).

**Fig. 5.**
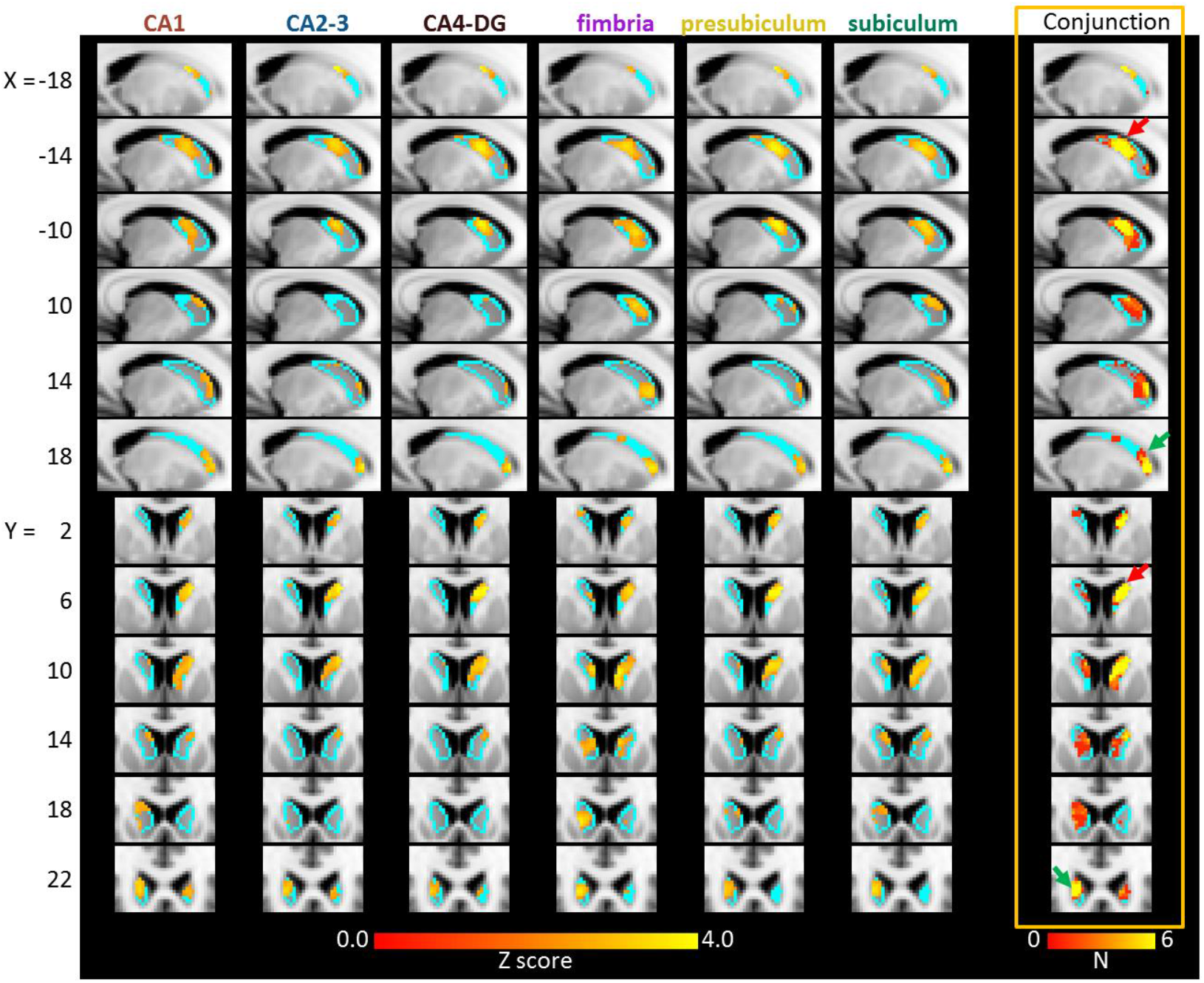
Association between navigation ability and seed-based functional connectivity in the caudate with each hippocampal subfield. The first six columns show the association results (p < 0.05, FWE-corrected) with CA1, CA2-3, CA4-DG, fimbria, presubiculum and subiculum, respectively. The last column shows the conjunction map of these associations.

Taken together, these findings showed a considerable navigation-specific nature of the association between the hippocampal-caudate interaction and spatial navigation, suggesting a critical role of this intrinsic interaction in supporting people’s navigation.

### Hippocampal-caudate interaction predicts pointing performance in the 3D pointing task

Based on these findings, we asked whether the hippocampal-caudate interactions would predict individual’s performance in a behavioral navigation task. Here, we used a pointing task in a virtual environment (Fig. 6A, 6B). As expected, the pointing error showed significant correlation with self-reported navigation ability (Fig. 6C: *r* = −0.22, *p* = 0.010). Similarly, males showed significantly smaller pointing error than female (t(158) = −5.14*,p* < 0.001). To rule out the possible influence of video game playing, participants were also asked to rate the frequency with which they played the video games requiring navigation through an environment. The mean self-rating of experience with video games was significantly lower for women (M = 1.47, SD = 0.77) than for men (M = 2.41, SD = 1.02), t(154) = 6.48, *p* < 0.001. A simple correlation analysis showed that higher video game experience was associated with lower overall pointing error in the experimental mazes, *r* = - 0.31, *p* < 0.001. Thus, video game experience, as well as sex and age, were controlled in the following analyses. Initially, we conducted correlation analyses between hippocampal-caudate interaction and pointing error, and found significant correlations in both ipsilateral and contralateral hippocampal-caudate interaction (Fig. 7) (HippL-CaudL: *r* = −0.20, *p* = 0.010; HippR-CaudR: *r* = −0.15, *p* = 0.039; HippL-CaudR: *r* = −0.18, *p* = 0.017; HippR-CaudL: *r* = −0.22, *p* = 0.005), which replicated our findings above. Moreover, we used SVR and LOOCV procedure to predict individual’s performance in the 3D pointing task based on these hippocampal-caudate interactions. As is shown in Fig. 8, the correlation between predicted and actual scores was 0.38 *(p* < 0.001), indicating that individual’s navigation performance could be well predicted using the measures of hippocampal-caudate interaction at rest.

**Fig. 6.**
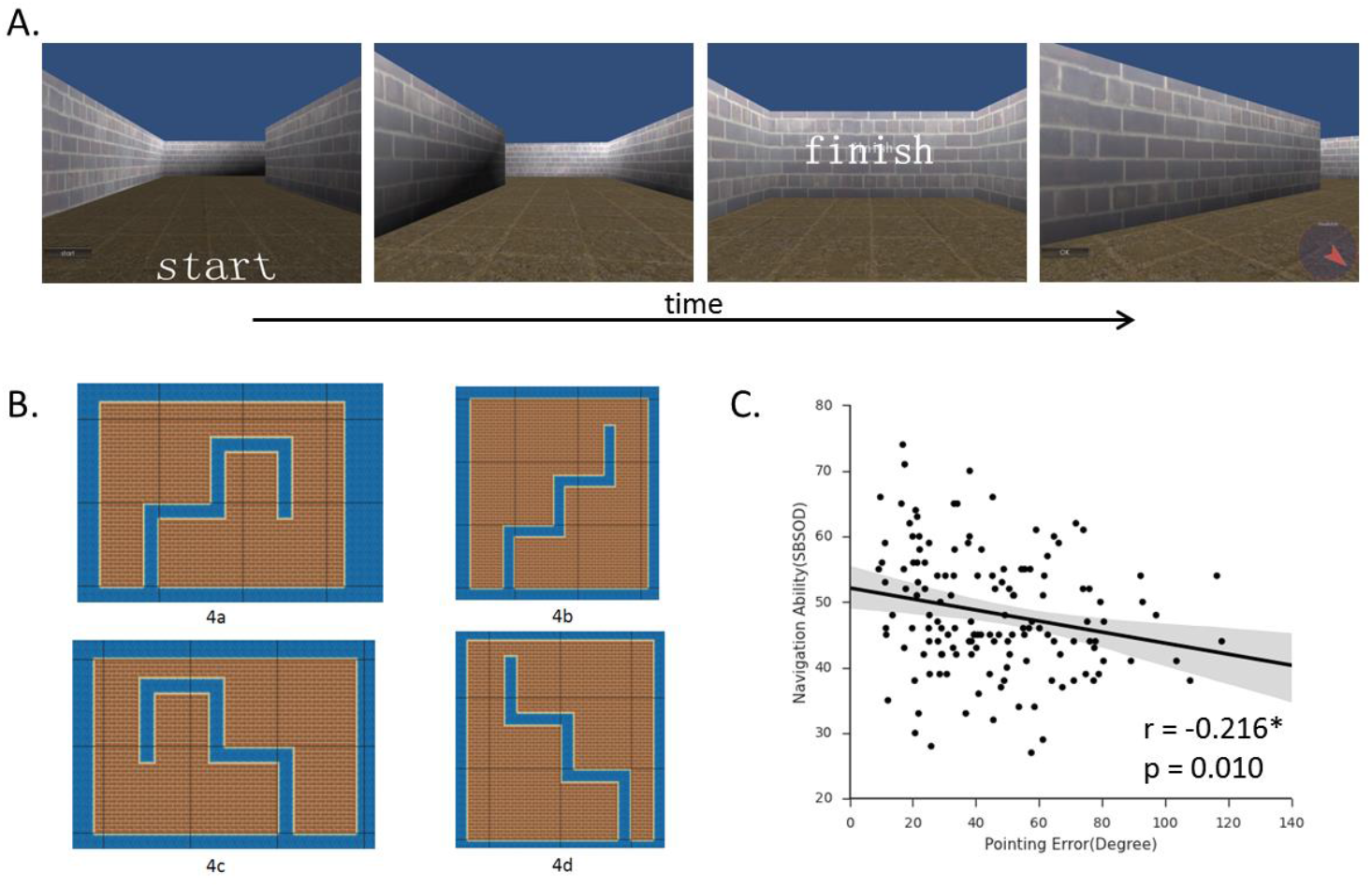
The 3D pointing task. (A) Screenshots from the task in the virtual environment, and (B) exemplar routes used in this study. (C) Correlation between pointing error in this task and navigation ability measured with SBSOD. Shaded regions depict 95% confidence intervals. * p < 0.05.

**Fig. 7.**
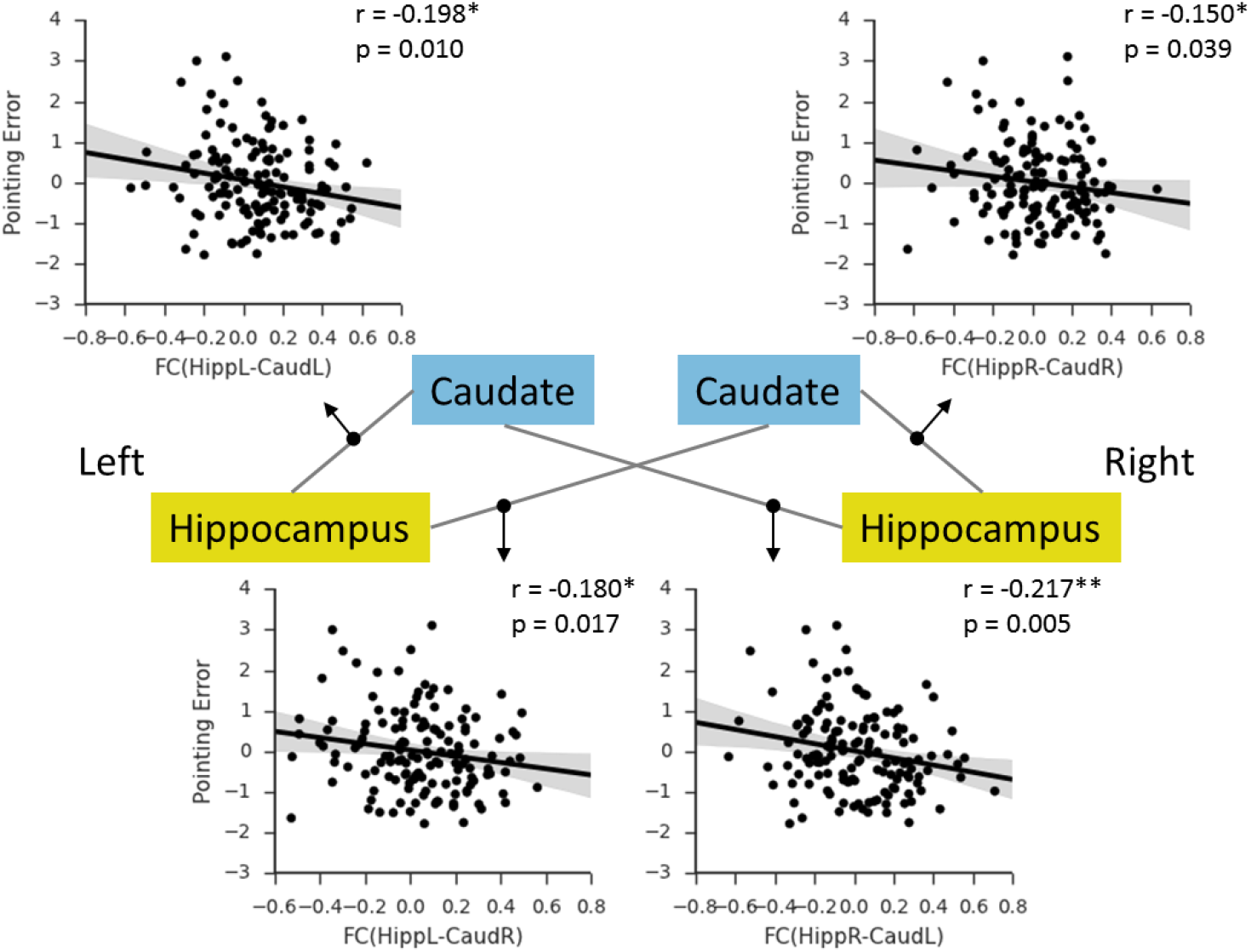
Correlations between the hippocampal-caudate functional connectivity and pointing error measured with 3D pointing task. Shaded regions depict 95% confidence intervals. HippR-CaudR/ HippL-CaudL, functional connectivity between the hippocampus and caudate in the same hemispheres; HippR-CaudL/ HippL_CaudR, functional connectivity between the hippocampus and caudate in the different hemispheres; R, right; L, left. * p < 0.05; ** p < 0.01

**Fig. 8.**
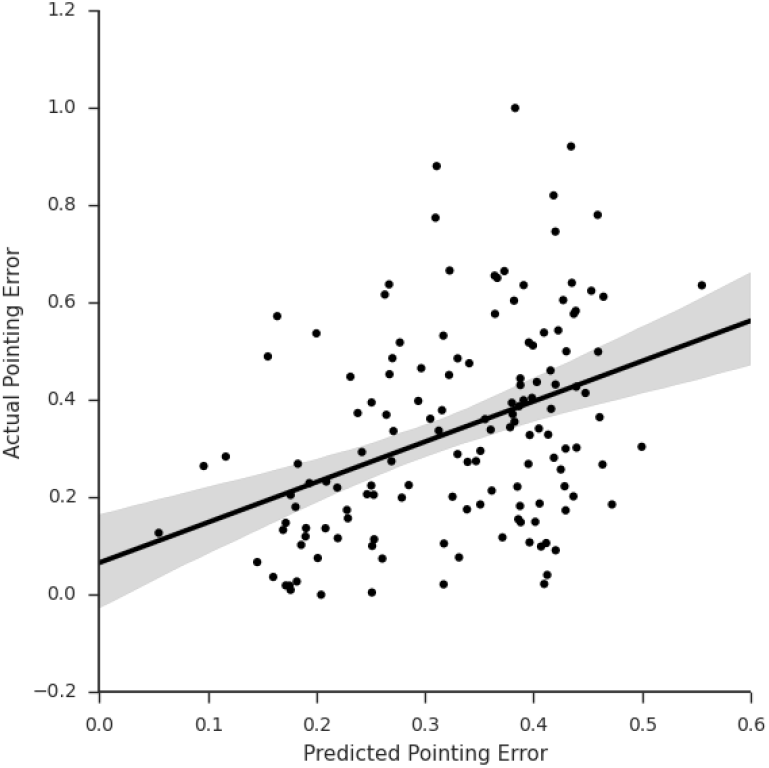
Scatterplot represents the linear association between predicted pointing error and actual pointing error. Shaded regions depict 95% confidence intervals. *** **p** < 0.001.

## Discussion

In sum, we investigated the functional interaction between the hippocampus and the caudate with resting-state fMRI, and confirmed its behavioral importance in human navigation. Specifically, we found slightly positive functional connectivity between the hippocampus and the caudate (in particular in good navigators), suggesting a cooperative nature of the interaction. Moreover, we found that the strength of the hippocampal-caudate interaction was positively correlated with people’s daily navigation performance measured with SBSOD. In addition, the observed brain-behavior association could not be accounted for by non-navigation abilities (e.g., general ability and small-scale spatial ability), which suggests a navigation-specific nature. Finally, we found that the observed associations could be replicated with a 3D pointing task in virtual reality, and more importantly, prediction analysis with machine learning algorithm and a cross-validation procedure showed that the specific navigation performance for each individual could be well predicted using the hippocampal-caudate interaction measures. Our findings support the hypothesis that in human the two distinct representation systems, the allocentric and egocentric systems, interact in a cooperative manner to support flexible navigation.

Our findings advanced previous studies linking the two spatial representation systems to spatial navigation. As a complex cognitive process, flexible navigation relies on proper coupling between multiple brain regions including the hippocampus and the caudate.

Though these two systems function differently in navigation (Morris RG *et al.* 1982; Cook D and RP Kesner 1988) as discussed in Introduction, the cooperative interaction between them has been strongly highlighted for its behavioral significance during navigation tasks (Voermans NC *et al.* 2004; Brown TI *et al.* 2012; Rice JP *et al.* 2015). For example, in patients with Huntington’s disease, the hippocampus compensates for gradual caudate dysfunction with a gradual activity increase during route recognition to maintaining normal behavior (Voermans NC *et al.* 2004). Our results advanced our understanding of the hippocampal-caudate interaction in the following senses. First, with a large dataset of resting-state fMRI, we investigated the intrinsic interaction between the hippocampus and the caudate without any task modulation. To the best of our knowledge, this study is the first report of a positive interaction between the hippocampus and the caudate at rest, which suggested intrinsic cooperative dynamics of two systems underlying navigation. Second, the functional interaction was further supported by the findings that the hippocampal-caudate functional connectivity varies as a function of both daily navigation performance (measured by the SBSOD) and specific navigation performance (measured by the 3D pointing task), showing the behavioral importance in supporting human navigation of the hippocampal-caudate interaction. Finally, our findings that the intrinsic interaction of interest predicted each individual’s performance in the out-of-scanner pointing task, suggest that the hippocampal-caudate interaction would be intrinsically involved in shaping human navigation behaviors. It seems that the hippocampal-caudate interaction not only benefits human navigation during navigation (Brown TI *et al.* 2012), but also during rest cooperatively prepares for upcoming navigation tasks. Further studies are needed to fully understand their roles in spatial navigation.

There are also studies suggesting a competitive interaction between the hippocampus and caudate, one or the other being optimal for specific tasks. For example, with task fMRI, a previous study has identified a caudate region whose activity was negatively correlation with the medial temporal regions, including the hippocampus, suggesting a negative relationship between these structures (Poldrack RA et al. 2001). In addition, a recent structural study has showed that the gray matter in the hippocampus was negatively correlated to the gray matter in the caudate nucleus, suggesting a competitive interaction between these two brain areas (Bohbot VD et al. 2007). There is also evidence from animal studies (Eichenbaum H et al. 1988; Packard MG 1999) showing that the hippocampus and caudate acquire different types of information during learning: the hippocampus appears to acquire spatial relationships whereas the caudate acquires stimulus-response associations. This proposed competition between hippocampus and caudate may reflect an adaptive mechanism for optimizing behavior depending upon task demands (Poldrack RA *et al.* 2001). It is important to note that this does not contradict our findings. Unlike these studies, the present study examined the hippocampal-caudate interactions with individual resting-state fMRI data, which allows us to explore inter-regional interactions within each individual and independent to any specific task. Human navigation is a complex behavior and engages interactions of multiple processes. Thus, together with the different roles of these two systems in navigation (Hartley T *et al.* 2003; Doeller CF *et al.* 2008), our findings suggest that the cooperative interaction between hippocampus and caudate may reflect the intrinsic flexibility to switch navigation strategies according to the external environment which would do benefit for accurate navigation (e.g., (Iglói K et al. 2009)). This conjecture is largely supported by the positive functional connectivity between the hippocampus and caudate (in particular in good navigators), and the positive association between the intrinsic hippocampal-caudate interaction and navigation performance. Our findings also coincide with previous neurophysiological evidence (Voermans NC *et al.* 2004) that the hippocampus compensates for the functional degradation of the caudate to maintain normal behavior in route recognition. Taken together, our findings provide novel evidence for cooperative contribution of the hippocampal-caudate interaction to human navigation. Note that the strength of the functional connectivity of interest was relatively weak (~ 0.10).

This finding is consistent to one recent study (Nyberg L et al. 2016), and more importantly, we showed that the weak connections could reliably predict individual differences in behavioral performance, suggesting a critical functional role of weak connections in human brain. Previous brain network studies have generally focused on strong connectivity patterns (e.g., top 5% connections or connections larger than 0.20), for both diagnostic purposes (Bassett DS et al. 2008; Wang J et al. 2015), and the understanding of the organization (Sporns O and JD Zwi 2004; Achard S et al. 2006) and developmental trajectories (Meunier D et al. 2009; Cao M et al. 2014) of human brain. In most of previous studies, weak connections are usually considered spurious and assigned a value of zero, resulting in the fact that the role of weak connections has remained obscure for years. This is somehow surprising, given that the relevance of weak connections in other complex systems had already been stressed many years ago. For instance, Granovertter (1973) has pointed out the importance of weak relationships (edges) between members (nodes) in complex social networks, for the maintenance of overall system dynamics, and the spreading of new ideas and information (Granovetter M 1973). Only recently, new evidence has suggested that weak connections may be a useful marker for general cognitive functioning (Santarnecchi E et al. 2014) and specific pathological conditions (Bassett DS et al. 2012). Our findings provide strong support for the hypothesis that weak connectivity reflects meaningful individual differences rather than simply a technical artifact, which could have substantial implications for functional connectivity study. This could also put forward the need to characterize the functional properties and dynamics of both weak and strong connections in order to obtain a comprehensive understanding of their role in considerable individual variability.

The present study provided important insight into the neural correlates of the considerable individual difference in navigation ability by linking the hippocampal-caudate interactions and spatial navigation. However, note that the exact mechanism underlying the observed hippocampus-caudate interaction during rest and its role in human spatial navigation is not clear yet. Previous studies suggest that the hippocampus and caudate do not share direct anatomical connections, but the medial prefrontal cortex (MPFC) receives direct projections from the hippocampus and sends direct projections to the medial caudate nucleus (Cavada C et al. 2000; Haber SN et al. 2006; Roberts AC et al. 2007). A recent study suggests that MPFC might internally simulate alternative strategies (Schuck NW et al. 2015), and in navigation researches, the MPFC has been suggested as a coordinator between striatal and hippocampal systems in rodents (Killcross S and E Coutureau 2003) and humans (Doeller CF *et al.* 2008). Further studies are needed to reveal the exact role of the MPFC in the hippocampus-caudate interactions and their roles in spatial navigation. The observed brain-behavior association might reflects efficiency of a more distributed brain network including the MPFC and further studies are needed to examine this possibility (Kong XZ, X Wang, *et al.* 2017). In addition, both genetic and environmental factors could contribute to individual differences in the hippocampal-caudate interaction and navigation. On the one hand, genetic factors predispose individual differences in the spontaneous brain function (Glahn DC et al. 2010) including the hippocampal-caudate interaction (Nyberg L *et al.* 2016) and in the task-evoked activity related to human navigation (Kong XZ, Y Song, *et al.* 2017). On the other hand, environmental factors such as acquired experience may be critical to reinforce these differences (Maguire EA et al. 2000; Woolley DG et al. 2015). Better navigation ability may reflect more navigation experience, which may boost the functional synchronization between the hippocampus and the caudate. This could be supported by greater functional connectivity between the hippocampus and the caudate revealed when participant uses contextual information to aim navigation (Brown TI *et al.* 2012). Finally, regardless of the mechanisms involved, our results indicate that the two memory systems interact cooperatively during rest, and that this interaction shapes individual’s navigation ability. The exact underlying physiological, genetic, and developmental mechanisms of the relation will be an important topic for further studies.

In sum, our findings provide first evidence for cooperative interaction of the hippocampus and caudate during rest, and its critical role in human navigation. Further research is needed to examine the dynamics of this interaction in response to different spatial tasks (e.g., spatial encoding and retrieval) and early development. Moreover, spatial navigation impairment is associated with both normal aging (Gazova I et al. 2012) and different psychiatric disorders, such as mild cognitive impairment (MCI) (Delpolyi AR et al. 2007; Hort J et al. 2007) and Alzheimer’s disease (AD) (Lithfous S et al. 2013). Previous studies have linked the deficit to some structural and/or functional alteration of the hippocampus and caudate (Rombouts SA et al. 2000; Chetelat G et al. 2002; Dai W et al. 2009). Systematic studies on the dynamic interaction between related structures would help us identify the previously ignored impact from altered inter-regional interaction, thus advance our understanding of the neural mechanisms underlying normal aging and psychiatric disorders. Finally, given the complex nature of human navigation, functional connectivity analysis of whole brain and/or multiple navigation-related regions would lead to better understanding of the dynamic interactions between multiple systems in spatial navigation.

## Acknowledgments

This study was funded by the National Natural Science Foundation of China (31230031), the National Basic Research Program of China (2014CB846101), the National Natural Science Foundation of China (31221003, 31471067 and 31470055), the National Social Science Foundation of China (13&ZD073, 14ZDB160), and Changjiang Scholars Programme of China. We thank Prof. Dr. Christian Doeller and Dr. Branka Milivojevic for helpful suggestions on the manuscript.

## Author Contributions

X.K., Y.P. and J.L. conceived and designed the experiments, X.K., X.W, and Z.Z conducted the experiments, X.K., Y.P. and X.W. analyzed the data, X.K., Y.P., S.X., X.H., and J.L. wrote the paper.

